# MolRep: A Deep Representation Learning Library for Molecular Property Prediction

**DOI:** 10.1101/2021.01.13.426489

**Authors:** Jiahua Rao, Shuangjia Zheng, Ying Song, Jianwen Chen, Chengtao Li, Jiancong Xie, Hui Yang, Hongming Chen, Yuedong Yang

**Affiliations:** School of Computer Science and Engineering, Sun Yat-Sen University; Galixir Technology; Center of Chemistry and Chemical Biology, Guangzhou Regenerative Medicine and Health Guangdong Laboratory

## Abstract

**Summary:** Recently, novel representation learning algorithms have shown potential for predicting molecular properties. However, unified frameworks have not yet emerged for fairly measuring algorithmic progress, and experimental procedures of different representation models often lack rigorousness and are hardly reproducible. Herein, we have developed MolRep by unifying 16 state-of-the-art models across 4 popular molecular representations for application and comparison. Furthermore, we ran more than 12.5 million experiments to optimize hyperparameters for each method on 12 common benchmark data sets. As a result, CMPNN achieves the best results ranked the 1st in 5 out of 12 tasks with an average rank of 1.75. Relatively, ECC has good performance in classification tasks and MAT good for regression (both ranked 1st for 3 tasks) with an average rank of 2.71 and 2.6, respectively.

**Availability:** The source code is available at: https://github.com/biomed-AI/MolRep

**Supplementary information:** Supplementary data are available online.

## 1 Introduction

The accurate characterization of molecule properties is one of the key tasks in drug development. Decades of computational modeling methods, such as quantitative structure–activity and structure–property relationships (QSAR/PR), provide alternative approaches to profile molecules and save a huge amount of costs and labor involved in experiments (Hecht and Fogel, 2009; Cherkasov *et al*., 2014). In recent years, with the substantial increase in available experimental molecular properties data points, there is growing interest in developing powerful machine learning methods especially deep learning methods for molecular property prediction, thereby accelerating the speed of drug development (Duvenaud *et al*., 2015; Gilmer *et al*., 2017; Yang *et al*., 2019b; Song *et al*., 2020).

Advance in these algorithms is both measured and steered by well-established molecular property benchmarks, such as the MoleculeNet (Wu *et al*., 2018) and the admetSAR (Yang *et al*., 2019a). Albeit promising results have been frequently reported, there are raised concerns about several deficiencies in scholarship, such as miss of strong baseline models, poor experimental reproducibility and replicability as well as inconsistent data splitting methods and performance metrics. Furthermore, recent property prediction studies were often lacking hyper-parameter optimization, and most models are not carefully tuned to achieve peak performance (Shen and Nicolaou, 2020). As a result, it can be difficult to assess the effectiveness of one method against another uniformly. These issues are not easy to settle, as extensive efforts are required to avoid improper practices. Comprehensive benchmark experiments across recently proposed deep representation methods will benefit the real performance improvement for any new modeling methodologies.

For this sake, we present MolRep: a Python package for training and evaluating deep representation learning models on chemical property datasets that provides uniform and rigorous comparisons over 16 state-of-the-art models derived from four different kinds of molecular representations including fixed descriptors/fingerprints, self-/unsupervised feature vectors, SMILES strings, and molecular graphs. We strictly followed the standard two-phase evaluation strategy (Cawley and Talbot, 2010), namely model selection on the valid set and assessment on the independent test set, avoiding over-optimistic and biased estimates of the true performance of a model. An elaborate hyper-parameter searching module was assembled to explore the best performance for each model.

Although there exist other toolkits like DeepChem (Wu *et al*., 2018) and admetSAR (Yang *et al*., 2019a), they either contained a few deep representation learning models or require many efforts to perform unified evaluation process and hyper-parameter searching. To the best of our knowledge, MolRep is the first framework specifically designed to facilitate the use of the latest deep representation learning models for users in the pharmaceutical community.

## 2 Software architecture

The MolRep framework consists of two modules: (i) the data preprocessing module and (ii) the model selection and assessment module(see Fig.1).

**Fig. 1.**
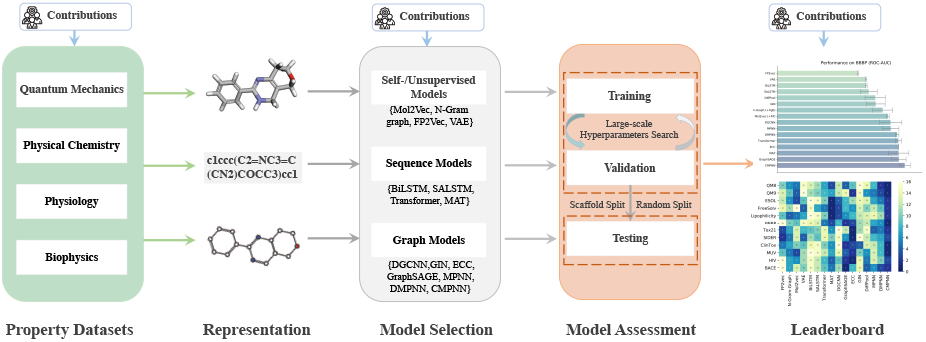
An overview of MolRep library.The open-source library currently provides 16 state-of-the-art deep representation learning models for fairly measuring algorithmic progress on molecular property prediction tasks.

### 2.1 Data preprocessing module

MolRep uses a unified data preprocessing pipeline that guarantees consistency in evaluating the performance of different models. Each molecule is then represented by SMILES string, molecular graphs, or three-dimensional conformer with RDKit. MolRep provided 16 initial molecule properties datasets from MoleculeNet and several literature, including Quantum Mechanics, Physical Chemistry, Biophysics, Physiology. Supplementary Table S1 lists details of these datasets.

### 2.2 Model selection and assessment module

MolRep has set up a standardized and uniform assessment module for molecular representation, such that the future models and datasets can be added to the existing modules for fair and objective comparison. Currently, we provided implementations of 16 competitive models that could be divided into self-/unsupervised learning models, sequence-based models, and graph-based models. The models were benchmarked on 12 datasets, and the comprehensive assessment includes a totally of 12.5 million experiments. The detailed information of these models is summarized in Supplementary Appendix A.

Since the performances of learning models are highly dependent on the particular test-set of choice, MolRep follows the rigorous assessment strategy of (Errica *et al*., 2019) by evaluating models on multiple test splits to achieve a fair comparison. For the sake of conciseness, the detailed assessment procedure is summarized in Supplementary Appendix B.

## 3 MolRep with CLI and Jupyter interface

MolRep has developed user-friendly APIs for uploading users’ own datasets in SDF or CSV format, updating the future learning models, and defining users’ own metrics such as Coefficient of determination (R-squared) or Pearson correlation coefficient. In the model assessment module, MolRep also provides options to add/modify/select hypermeters of each model. Additionally, this package is accessible and facile for chemists without expert knowledge in machine learning through an interactive Jupyter and command line (CLI) interface.

## 4 Results and Discussion

MolRep provides a complete re-evaluation of 16 state-of-the-art deep representation models over 16 kinds of molecular properties, which requires a significant amount of time and computational resources. The uniform package helps us fairly compare with the state-of-the-art representation models.

By comparing the results, we found that no single model or molecular representation performs consistently well on all tasks, suggesting that the development of deep representation learning models for properties prediction has not been exploited yet. Experimental results on 12 common benchmark data sets are detailed in Table S11-12.

Overall, CMPNN (Song *et al*., 2020) achieves the best results ranked the 1st in 5 out of 12 tasks with an average rank of 1.75. Relatively, ECC (Shang *et al*., 2018) has good performance in classification tasks and MAT (Maziarka *et al*., 2020) good for regression (both ranked 1st for 3 tasks) with an average rank of 2.71 and 2.6, respectively. Supplementary Figure 2 reports the overall ranks of all models on all datasets. Future ensemble models can be easily developed based on the presented leaderboard.

Finally, we provided the property prediction community with reliable and reproducible results to which researchers can compare their architectures. We hope that this work, along with the framework we release, will prove useful for practitioners to compare QSAR/PR models in a more rigorous way.

