## Supplemental information for "MolRep: A Deep Representation Learning Library for Molecular Property Prediction"

### Outline

#### Appendix A. Further details on MolRep.

1. **Overview of MolRep framework.** MolRep is a Python package for training and evaluating deep representation learning models on chemical property datasets. In particular, MolRep provides uniform and rigorous comparisons over 16 state-of-the-art models derived from four different kind of molecular representations including fixed descriptors/fingerprints, unsupervised feature vector, SMILES string and molecular graph.

2. **Molecular representations learning models implemented in MolRep.** MolRep provides 16 state-of-the-art models, including 4 unsupervised-learning models, 4 sequenced-learning models and 8 graph-learning models.

**Self-/Unsupervised Models.**

1. Mol2Vec[1] is an unsupervised approach to learns vector representations of molecular substructures that point in similar directions for chemically related substructures.
2. N-gram graph[2] is a simple unsupervised representation for molecules that first embeds the vertices in the molecule graph and then constructs a compact representation for the graph by assembling the vertex embeddings in short walks in the graph.
3. FP2Vec[3] is a molecular featurizer that represents a chemical compound as a set of trainable embedding vectors and combine with CNN model.
4. VAE[4] is a framework for training two neural networks (encoder and decoder) to learn a mapping from high-dimensional molecular representation into a lower-dimensional space.

**Sequence Models.**

1. BiLSTM[5] is an artificial recurrent neural network (RNN) architecture to encoding sequences from compound SMILES strings.
2. SALSTM[6] is a self-attention mechanism with improved BiLSTM for molecule representation.
3. Transformer[7] is a network based solely on attention mechanisms and dispensing with recurrence and convolutions entirely to encodes compound SMILES strings.
4. MAT[8] is a molecule attention transformer utilized inter-atomic distances and the molecular graph structure to augment the attention mechanism.

**Graph Models.**

1. DGCNN[9] is a deep graph convolutional neural network that proposes a graph convolution model with SortPooling layer which sorts graph vertices in a consistent order to learning the embedding of molecular graph.
2. GraphSAGE[10] is a framework for inductive representation learning on molecular graphs that used to generate low-dimensional representations for atoms and performs sum, mean or max-pooling neighborhood aggregation to updates the atom representation and molecular representation.
3. GIN[11] is the Graph Isomorphism Network that builds upon the limitations of GraphSAGE to capture different graph structures with the Weisfeiler-Lehman graph isomorphism test.
4. ECC[12] is an Edge-Conditioned Convolution Network that learns a different parameter for each edge label (bond type) on the molecular graph, and neighbor aggregation is weighted according to specific edge parameters.

5. DiffPool[13] combines a differentiable graph encoder with its an adaptive pooling mechanism that collapses nodes on the basis of a supervised criterion to learning the representation of molecular graphs.
6. MPNN[14] is a message-passing graph neural network that learns the representation of compound molecular graph. It mainly focused on obtaining effective vertices (atoms) embedding.
7. DMPNN[15] is another message-passing graph neural network that messages associated with directed edges (bonds) rather than those with vertices. It can make use of the bond attributes.
8. CMPNN[16] is the graph neural network that improve the molecular graph embedding by strengthening the message interactions between edges (bonds) and nodes (atoms).

3. **Datasets included in MolRep.**MolRep currently includes over 300,000 compounds with various properties. These properties could be divided into four categories: Quantum Mechanics (QM7, QM8, QM9), Physical Chemistry (ESOL, FreeSolv, Lipophilicity), Physiology (BBBP, Tox21, SIDER, ClinTox, hERG, Liver injury, Mutagenesis), and Biophysics (MUV, HIV, BACE). The summary of the 16 datasets can be found in Supplementary Table 1.

#### Appendix B. More details of the Assessment Module.

In model assessment module, the whole dataset was randomly split into K folds to form K test splits. For each test split, the remaining data was used to select optimal hyper-parameters through a standard n-fold cross-validation strategy, and the optimal hyperparameters were used to train a model on the whole training set for test on the test split. The obtained K results are then averaged to provide a stable estimate of the model performance. Here, we set both n and K as five. The evaluation metrics is dataset-specific according to previous studies, including the Area Under the Receiver Operating Characteristics (ROC-AUC) curve and the Precision-Recall Curve (PRC-AUC) for classification tasks, and Root Mean Square Error (RMSE) and mean absolute error (MAE) for regression tasks

The hyper-parameters was searched via the grid searching strategy. MolRep has compiled unified hyperparameters for each type of models that also include the hyper-parameters used by the authors in their respective papers. The number of hyperparameter combinations of all models is detailed in Supplementary Table S2. A full list of hyper-parameters can be found in Supplementary Table S3-10.

**Supplementary Table 1. Summary of the 16 Molecule Properties datasets included MolRep.**

| Category | Dataset | Task | Task type | #Molecule | Splits | Metric | Source |
|---|---|---|---|---|---|---|---|
| Quantum Mechanics | QM7 | 1 | Regression | 7160 | Stratified | MAE | Wu et al.[17] |
| | QM8 | 12 | Regression | 21786 | Random | MAE | Wu et al.[17] |
| | QM9 | 12 | Regression | 133885 | Random | MAE | Wu et al.[17] |
| Physical Chemistry | ESOL | 1 | Regression | 1128 | Random | RMSE | Wu et al.[17] |
| | FreeSolv | 1 | Regression | 642 | Random | RMSE | Wu et al.[17] |
| | Lipophilicity | 1 | Regression | 4200 | Random | RMSE | Wu et al.[17] |
| Physiology | BBBP | 1 | Classification | 2039 | Scaffold | ROC-AUC | Wu et al.[17] |
| | Tox21 | 12 | Classification | 7831 | Random | ROC-AUC | Wu et al.[17] |
| | SIDER | 27 | Classification | 1427 | Random | ROC-AUC | Wu et al.[17] |
| | ClinTox | 2 | Classification | 1478 | Random | ROC-AUC | Wu et al.[17] |
| | Liver injury | 1 | Classification | 2788 | Random | ROC-AUC | Xu et al.[18] |
| | Mutagenesis | 1 | Classification | 6511 | Random | ROC-AUC | Hansen et al.[19] |
| | hERG | 1 | Classification | 4813 | Random | ROC-AUC | Li et al.[20] |
| Biophysics | MUV | 17 | Classification | 93087 | Random | PRC-AUC | Wu et al.[17] |
| | HIV | 1 | Classification | 41127 | Scaffold | ROC-AUC | Wu et al.[17] |
| | BACE | 1 | Classification | 1513 | Scaffold | ROC-AUC | Wu et al.[17] |

**Supplementary Table 2. Hyperparameters Combinations of all models**

| Category | Models | Hyperparameters Combinations |
|---|---|---|
| Self/unsupervised Models | Mol2Vec | 720 |
| | N-Gram graph | 648 |
| | FP2Vec | 1296 |
| | VAE | 5184 |
| Sequence Models | BiLSTM | 1152 |
| | SALSTM | 3456 |
| | Transformer | 2592 |
| | MAT | 3456 |
| Graph Models | DGCNN | 3456 |
| | GraphSAGE | 1944 |
| | GIN | 3456 |
| | ECC | 3456 |
| | DiffPool | 3888 |
| | MPNN | 3072 |
| | DMPNN | 1944 |
| | CMPNN | 1944 |

#### Supplementary Table 3. N-Gram graph (Xgb) and Mol2Vec (RF) hyperparameter ranges

| | N-Gram graph (Xgb) | | Mol2Vec |
|---|---|---|---|
| booster | gbtree | Number-of-Trees | 100, 400, 1000 |
| Max-depth | 5, 10, 50, 100 | Max-features | None, sqrt, log2 |
| Learning-rate | 1, 3e-1, 1e-1, 3e-2 | Min-samples-leaf | 1, 10, 100, 1000 |
| n-estimators | 30, 100, 300, 1000, 3000 | Min-samples-leaf | 1, 3 |
| Min-child-weight | 1, 5, 10 | Class-weight | None, balanced_subsample, balanced |
| Max-delta-step | 0, 10, 20 | Max-leaf-nodes | None, 100, 500 |

#### Supplementary Table 4. FP2Vec hyperparameter ranges

| | FP2Vec |
|---|---|
| Batch-size | 32, 64, 128 |
| Learning-rate | 0.01, 0.001, 0.0001 |
| Epochs | 80, 100, 200 |
| Optimizer | Adam |
| Hidden-size | 128, 256, 512, 1024 |
| Window-size | 5, 8, 16 |
| Bit-size | 1024, 2048 |
| Weight-decay | 0.0, 0.1 |

#### Supplementary Table 5. VAE hyperparameter ranges

| | VAE |
|---|---|
| Batch-size | 32, 64, 128 |
| Learning-rate | 0.01, 0.001, 0.0001 |
| Epochs | 30, 100 |
| Optimizer | Adam |
| Encoder-hidden-size | 64, 256 |
| Encoder-layers | 2, 4, 8 |
| Encoder-dropout | 0.2, 0.5 |
| Z-embedding | 64, 256 |
| Decoder-hidden-size | 128, 256, 512 |
| Decoder-layers | 2, 3 |
| Decoder-dropout | 0.0, 0.1 |

#### Supplementary Table 6. BiLSTM and SALSTM hyperparameter ranges

| | BiLSTM | SALSTM |
|---|---|---|
| Batch-size | 16, 32, 64 | 16, 32, 64 |
| Learning-rate | 0.1, 0.01, 0.001 | 0.1, 0.01, 0.001 |
| Epochs | 30, 50, 100, 300 | 30, 50, 100, 300 |
| Hidden-size | 64, 128, 256, 512 | 64, 128, 256, 512 |
| Optimizer | Adam, SGD | Adam, SGD |
| Early-stopper | Patience(patience=500), Patience(patience=300) | Patience(patience=500) , Patience(patience=300) |
| Attention-hops | - | 5, 10, 15 |
| Weight-decay | 0.0, 0.001 | 0.0, 0.001 |

#### Supplementary Table 7. Transformer hyperparameter ranges

| | Transformer |
|---|---|
| Batch-size | 32, 64 |
| Learning-rate | 0.01, 0.0001 |
| Number-of-layers | 2, 6, 8 |
| Epochs | 10, 30, 100 |
| Hidden-size | 128, 256, 512 |
| Optimizer | Adam |
| Early-stopper | Patience(patience=500) |
| Dropout | 0.0, 0.2 |
| Attention-heads | 2, 4, 8 |
| Attention-dropout | 0.1, 0.2 |
| Intermediate-size | 128, 256 |

#### Supplementary Table 8. MAT hyperparameter ranges

| | MAT |
|---|---|
| Batch-size | 16, 32, 64 |
| Learning-rate | 0.01, 0.0001 |
| Number-of-layers | 4, 6, 8 |
| Epochs | 30, 100 |
| Hidden-size | 256, 512 |
| Optimizer | Adam |
| Early-stopper | Patience(patience=500) |
| Dropout | 0.0, 0.1 |
| Attention-heads | 1, 4, 8 |
| Attention-lambda | 0.1, 0.3 |
| Distance-kernel | 'Softmax', 'Exp' |
| Weight-decay | 0.0, 0.1 |

#### Supplementary Table 9. DGCNN, GraphSAGE, GIN, ECC, and DiffPool hyperparameter ranges

| | DGCNN | GraphSAGE | GIN | ECC | DiffPool |
|---|---|---|---|---|---|
| Batch-size | 32, 50, 64 | 32, 64, 128 | 32, 64, 128 | 32, 64, 128 | 16, 32 |
| Learning-rate | 0.01, 0.001, 0.0001 | 0.01, 0.001, 0.0001 | 0.01, 0.001 | 0.1, 0.01 | 0.001, 0.0001, 0.00001 |
| Epochs | 50, 100, 300 | 50, 100, 300 | 50, 100, 300 | 50, 100, 300 | 50, 100, 300 |
| Hidden-size | 32, 64, 128, 256 | 32, 64, 128, 256 | [64, 64, 64, 64], [32, 32, 32, 32], [32, 32, 64, 64], [64] | 32, 64, 128, 256 | 32, 64 |
| Number-of-layers | 2, 3, 4, 8 | 3, 5 | - | 1, 2, 4, 8 | 1, 2, 4 |
| Optimizer | Adam | Adam | Adam | Adam, SGD | Adam, SGD |
| Early-stopper | Patience | Patience | Patience | Patience | Patience |
| Dropout | - | - | 0.0, 0.5 | 0.05, 0.25, 0.1 | - |
| Weight-decay | 0.0, 0.1 | 0.0, 0.1, 0.2 | 0.0, 0.1, 0.2 | 0.0, 0.1 | 0.0, 0.1 |
| Others | Dense-dim:{64, 128}; K:{0.9, 0.6} | Aggregation: {‘add’, ‘max’, ‘mean’}; | Aggregation: {‘sum’, ‘mean’}; stepLR: {gramma: 0.3, 0.5; step_size: 10, 50} | - | GNN-dim-hidden: {32, 64, 128}; MLP-dim-hideen: {50, 64, 128} |

#### Supplementary Table 10. MPNN, DMPNN, and CMPNN hyperparameter ranges

| | MPNN | DMPNN | CMPNN |
|---|---|---|---|
| Batch-size | 32, 50, 64 | 32, 50, 64 | 32, 50, 64 |
| Learning-rate | 0.0001, 0.00001 | 0.0001, 0.00001 | 0.0001, 0.00001 |
| Epochs | 10, 30, 100, 300 | 10, 30, 500 | 10, 30 |
| Hidden-size | 256, 512 | 256, 300, 512 | 256, 300 |
| Depth | 1, 3, 4, 8 | 2, 3, 4 | 2, 3, 5 |
| Optimizer | Adam | Adam | Adam |
| Early-stopper | Patience | Patience | Patience |
| Dropout | 0.0, 0.2 | 0.0, 0.2 | 0.0, 0.1, 0.2 |
| FFN-hidden-size | 256, 300 | 256, 300 | 256, 300, 512 |
| FFN-num-layers | 2, 4 | 2, 3, 4 | 2, 3, 5 |
| Aggregation-norm | 64, 100 | - | - |

#### Supplementary Table 11. Performance on Regression datasets.

Comparisons of prediction performance between 16 state-of-the-art models on 5 regression datasets in terms of MAE or RMSE. Best performances are highlighted in bold.

| Datasets | **Quantum Mechanics** | | **Physical Chemistry** | | |
|---|---|---|---|---|---|
| | QM8 | QM9 | ESOL | FreeSolv | Lipophilicity |
| Methods | MAE | MAE | RMSE | RMSE | RMSE |
| FP2vec | 0.0236±0.0005 | 3.1688±0.1287 | 1.0653±0.0987 | 1.3479±0.2203 | 0.8781±0.0319 |
| N-Gram-Graph (+Xgb) | 0.0193±0.0001 | 2.9971±0.0099 | 0.6832±0.0180 | 1.2361±0.1183 | 0.7552±0.0261 |
| Mol2vec (+RF) | 0.0188±0.0000 | 2.7739±0.0067 | 0.6239±0.0039 | 1.4791±0.2015 | 0.7783±0.0014 |
| VAE | 0.0311±0.0001 | 3.3372±0.1992 | 0.8976±0.0583 | 0.9671±0.0958 | 0.7834±0.0798 |
| BiLSTM | 0.0299±0.0000 | 3.1103±0.0087 | 1.1391±0.1309 | 1.3498±0.2091 | 1.0356±0.0275 |
| SALSTM | 0.0278±0.0002 | 3.0572±0.1094 | 1.1213±0.0372 | 1.0387±0.1092 | 1.0098±0.0327 |
| Transformer | 0.0199±0.0000 | 2.9112±0.0197 | 0.9183±0.0397 | 0.9976±0.0075 | 0.9087±0.00648 |
| MAT | 0.0120±0.0001 | 2.8977±0.0009 | **0.3012±0.0244** | **0.3188±0.0003** | **0.4592±0.0342** |
| DGCNN | 0.0287±0.0003 | 3.1387±0.2108 | 0.6505±0.0301 | 0.9191±0.1267 | 0.7002±0.0325 |
| GraphSAGE | 0.0187±0.0001 | 3.1863±0.1037 | 0.8506±0.0191 | 1.0749±0.1073 | 0.7791±0.0105 |
| ECC | 0.0205±0.0004 | <u>2.5496±0.0329</u> | 0.9982±0.0073 | 1.0021±0.0582 | 0.8697±0.0031 |
| GIN | 0.0337±0.0005 | 3.2416±0.0840 | 1.1046±0.0000 | 1.3213±0.0000 | 0.9322±0.0000 |
| DiffPool | 0.0310±0.0000 | 2.8964±0.0003 | 1.0720±0.0032 | 1.2129±0.0301 | 0.9578±0.0013 |
| MPNN | 0.0202±0.0011 | 2.8178±0.0239 | 0.7021±0.0138 | 1.2421±0.2890 | 0.9872±0.0432 |
| DMPNN | <u>0.0118±0.0002</u> | 2.6823±0.0063 | 0.6623±0.0482 | 1.1572±0.1030 | 0.9234±0.0138 |
| CMPNN | **0.0108±0.0006** | **2.5090±0.0031** | <u>0.5508±0.0149</u> | <u>0.8134±0.0223</u> | <u>0.6128±0.0106</u> |

#### Supplementary Table 12. Performance on Classification datasets

Comparisons of prediction performance between 16 state-of-the-art models on 5 classification datasets in terms of ROC-AUC or PRC-AUC. Best performances are highlighted in bold.

| Datasets | Physiology | | | | Biophysics | | |
|---|---|---|---|---|---|---|---|
| | BBBP | Tox21 | SIDER | ClinTox | MUV | HIV | BACE |
| Methods | ROC-AUC | ROC-AUC | ROC-AUC | ROC-AUC | PRC-AUC | ROC-AUC | ROC-AUC |
| FP2vec | 0.8076±0.0032 | 0.8578±0.0076 | <u>0.6678±0.0068</u> | 0.8834±0.0432 | 0.0856±0.0031 | 0.7894±0.0052 | 0.8129±0.0492 |
| N-Gram-Graph (+Xgb) | 0.9012±0.0385 | 0.8371±0.0421 | 0.6482±0.0437 | 0.8753±0.0077 | 0.1011±0.0000 | 0.8378±0.0034 | 0.8472±0.0057 |
| Mol2vec (+RF) | 0.9213±0.0052 | 0.8139±0.0081 | 0.6043±0.0061 | 0.8572±0.0054 | 0.1178±0.0032 | 0.8413±0.0047 | 0.8284±0.0023 |
| VAE | 0.8378±0.0031 | 0.8315±0.0382 | 0.6493±0.0762 | 0.8674±0.0124 | 0.0794±0.0001 | 0.8109±0.0381 | 0.8368±0.0762 |
| BiLSTM | 0.8391±0.0032 | 0.8279±0.0098 | 0.6092±0.0303 | 0.8319±0.0120 | 0.0382±0.0000 | 0.7962±0.0098 | 0.8263±0.0031 |
| SALSTM | 0.8482±0.0329 | 0.8253±0.0031 | 0.6308±0.0036 | 0.8317±0.0003 | 0.0409±0.0000 | 0.8034±0.0128 | 0.8348±0.0019 |
| Transformer | 0.9610±0.0119 | 0.8129±0.0013 | 0.6017±0.0012 | 0.8572±0.0032 | 0.0716±0.0017 | 0.8372±0.0314 | 0.8407±0.0738 |
| MAT | 0.9620±0.0392 | 0.8393±0.0039 | 0.6276±0.0029 | 0.8777±0.0149 | 0.0913±0.0001 | 0.8653±0.0054 | 0.8519±0.0504 |
| DGCNN | 0.9311±0.0434 | 0.7992±0.0057 | 0.6007±0.0053 | 0.8302±0.0126 | 0.0438±0.0000 | 0.8297±0.0038 | 0.8361±0.0034 |
| GraphSAGE | <u>0.9630±0.0274</u> | 0.8166±0.0041 | 0.6403±0.0045 | <u>0.9116±0.0146</u> | 0.1145±0.0000 | <u>0.8705±0.0724</u> | **0.9316±0.0360** |
| ECC | 0.9620±0.0003 | **0.8677±0.0090** | **0.6750±0.0092** | 0.8862±0.0831 | <u>0.1308±0.0013</u> | **0.8733±0.0025** | 0.8419±0.0092 |
| GIN | 0.8746±0.0359 | 0.8178±0.0031 | 0.5904±0.0000 | 0.8842±0.0004 | 0.0832±0.0000 | 0.8015±0.0328 | 0.8275±0.0034 |
| DiffPool | 0.8732±0.0391 | 0.8012±0.0130 | 0.6087±0.0130 | 0.8345±0.0233 | 0.0934±0.0001 | 0.8452±0.0042 | 0.8592±0.0391 |
| MPNN | 0.9321±0.0312 | 0.8440±0.014 | 0.6313±0.0121 | 0.8414±0.0294 | 0.0572±0.0001 | 0.8032±0.0092 | 0.8493±0.0013 |
| DMPNN | 0.9562±0.0070 | 0.8429±0.0391 | 0.6378±0.0329 | 0.8692±0.0051 | 0.0867±0.0032 | 0.8137±0.0072 | 0.8678±0.0372 |
| CMPNN | **0.9854±0.0215** | <u>0.8593±0.0088</u> | 0.6581±0.0020 | **0.9169±0.0065** | **0.1435±0.0002** | 0.8687±0.0003 | <u>0.8932±0.0019</u> |

**Supplementary Figure 1. Comparisons of prediction performance between MAT, N-Gram graph and CMPNN.**

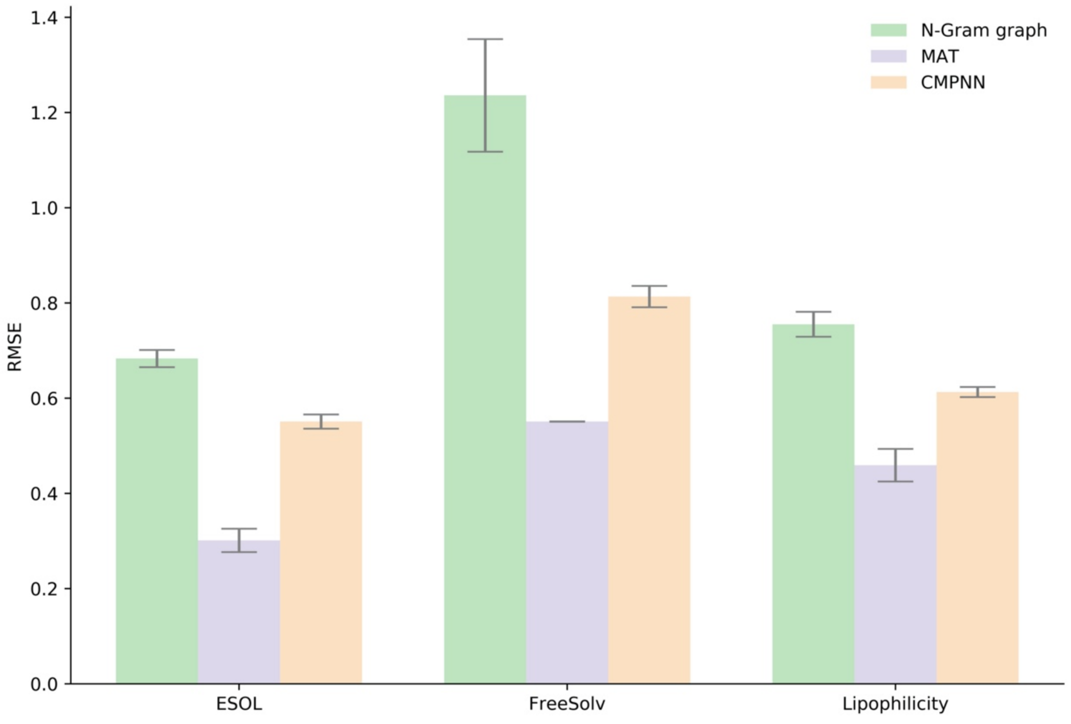

(a). Performance on regression datasets. (lower is better)

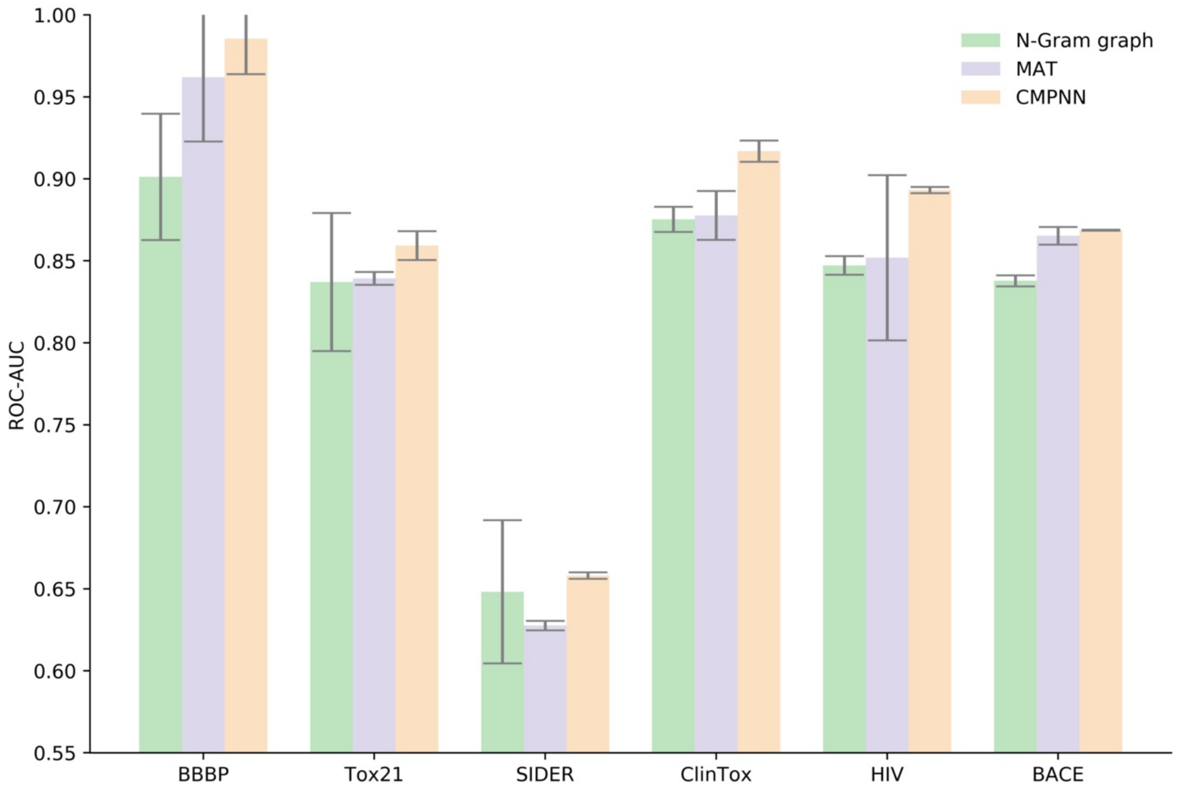

(b) Performance on classification datasets (higher is better)

**Supplementary figure 2. The overall ranks of all models on all datasets.**

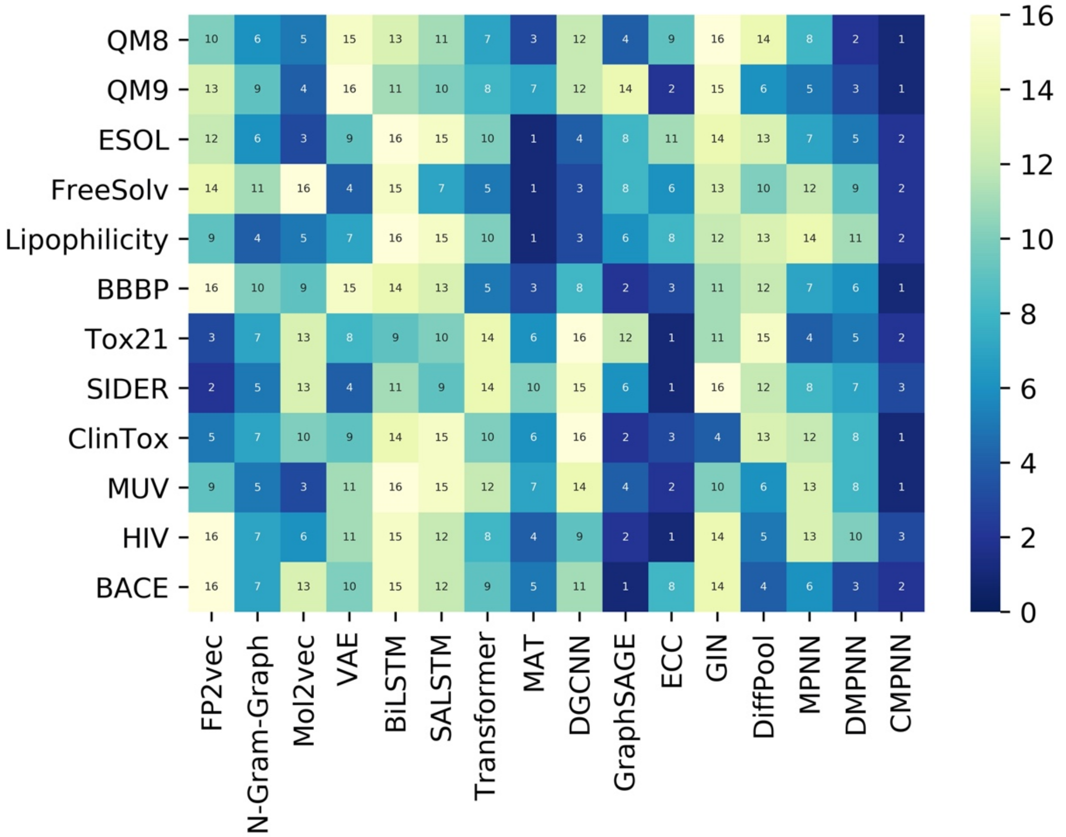
